# Immunologic changes are detectable in the peripheral blood transcriptome of clinically asymptomatic Chagas cardiomyopathy patients

**DOI:** 10.1101/2023.10.03.560680

**Authors:** Carolina Duque, Jaime So, Yagahira E. Castro-Sesquen, Kelly DeToy, Sneider A. Gutierrez Guarnizo, Fatemeh Jahanbakhsh, Edith Malaga Machaca, Monica Miranda-Schaeubinger, Indira Chakravarti, Virginia Cooper, Mary E. Schmidt, Luigi Adamo, Rachel Marcus, Kawsar R. Talaat, Robert H. Gilman, Monica R. Mugnier, the Chagas Working Group

## Abstract

Chagas disease, caused by the protozoan parasite *Trypanosoma cruzi*, is a neglected parasitic disease that affects approximately 6 million individuals worldwide. Of those infected, 20-30% will go on to develop chronic Chagas cardiomyopathy (CCC), and ultimately many of these individuals will progress to advanced heart failure. The mechanism by which this progression occurs is poorly understood, as few studies have focused on early CCC. In this study, we sought to understand the physiologic changes associated with *T. cruzi* infection and the development of CCC. We analyzed gene expression in the peripheral blood of asymptomatic Chagas patients with early structural heart disease, Chagas patients without any signs or symptoms of disease, and Chagas-negative patients with and without early structural heart disease. Our analysis shows that early CCC was associated with a downregulation of various peripheral immune response genes, with gene expression changes suggestive of reduced antigen presentation and T cell activation. Notably, these genes and processes were distinct from those of early cardiomyopathy in Chagas-negative patients, suggesting that the processes mediating CCC may be unique from those mediating progression to other cardiomyopathies. This work highlights the importance of the immune response in early CCC, providing insight into the early pathogenesis of this disease. The changes we have identified may serve as biomarkers of progression and could inform strategies for the treatment of CCC in its early stages, before significant cardiac damage has occurred.

## Introduction

Chagas disease, caused by the protozoan parasite *Trypanosoma cruzi*, is estimated to affect 6 million individuals worldwide, with an additional 70 million at risk [1, 2]. Of those infected, 20-30% will progress to chronic Chagas cardiomyopathy (CCC), a dilated cardiomyopathy resulting in symptoms of heart failure, cardiac arrhythmias, strokes, pulmonary embolisms, and/or sudden cardiac death, with significant morbidity and an estimated 12,000 deaths per year [1, 3, 4]. Histologically, advanced CCC is characterized by extensive fibrosis, necrosis, and inflammatory infiltrates [5-7]. Currently, there is no way to predict which patients will go on to develop CCC [8] and the mechanisms that underly disease progression are still poorly understood.

The natural history of Chagas cardiomyopathy is highly variable. Acute infection is most commonly very mild or asymptomatic and thus the disease is rarely diagnosed in the acute stage when anti-trypanosomal drugs are most effective[9-11]. Patients who are not treated in the acute phase will enter an indeterminate phase with no signs or symptoms of disease. While the majority of these patients will remain asymptomatic for their entire lives, approximately a third will gradually progress toward chronic cardiomyopathy decades after initial infection[4]. Alterations on electrocardiogram are one of the earliest clinical indications of CCC, with the most typical being right bundle branch blocks (RBBB), left anterior fascicular blocks (LAFB), and frequent premature ventricular contractions (PVC)[12, 13]. These electrical changes are a result of significant structural damage to the heart’s conduction system[14, 15], but the biologic processes contributing to this damage remain poorly understood.

Despite a dearth of research on the early processes involved in progression to CCC, evidence suggests the immune system plays an important role in the development of the cardiac damage that results in CCC. While *T. cruzi* is known to cause DNA damage, oxidative stress, and cell lysis[16], and thus is capable of directly damaging the heart, parasites can be difficult to detect in blood and tissues during the chronic stage of infection, thereby calling into question the extent to which the parasite alone is responsible for chronic tissue damage[17-21]. In line with this, a growing body of evidence has suggested that the host immune system plays a role in CCC pathogenesis[22]. Histologic studies indicate that CCC hearts, compared to other non-Chagas dilated cardiomyopathy hearts, have significantly increased CD8+ T cells, memory T cells[21, 23, 24], B cells[23], macrophages[23, 25], and mast cells[24] infiltrating cardiac tissue, all of which can contribute to tissue damage. In the peripheral blood, proinflammatory cells such as activated CD4+ T cells, NKT cells, cytotoxic NK cells[26], degranulating double-positive T cells[27], inflammatory monocytes[28], and cytokines such as TNF, INF-*γ*, and IL-6 are also increased[29].

While together these data suggest a role for the immune system in the development of CCC, most observations implicating the immune system in CCC have been derived from tissues acquired during late-stage, symptomatic disease [5, 6, 22, 30]. It thus remains unclear what role the immune system may play in the earliest stages of CCC and whether early immunologic changes could mediate and/or signal the development of CCC in a subset of infected patients. A better understanding of these early changes is essential given that early CCC is thought to be considerably more amenable to treatment and prevention than advanced CCC, where extensive cardiac damage has already occurred[8, 31].

In this study, we performed RNA sequencing analysis of peripheral blood samples from Chagas patients in the United States who originated from endemic regions. We focus on individuals with electrocardiographic alterations but without advanced symptomatic heart failure, with the goal of identifying very early changes that characterize CCC progression and that could serve as potential biomarkers of early CCC. We identified numerous immunologic transcriptomic changes that may help differentiate early CCC from non-Chagasic cardiomyopathy. Our findings suggest that the downregulation of certain immune system components may be an early indicator of CCC development.

## Methods

### Study design and participants

This study was performed under the ethics committee of the Johns Hopkins School of Public Health (protocol IRB 6713) and all patients provided written informed consent. From February 2016 to April 2018, patients originally from Chagas-endemic countries were recruited from health fairs and recruitment events held at various churches, community centers, and consulates, in Virginia, Maryland and Washington D.C[32, 33]. Patients provided a blood sample and underwent a cardiovascular evaluation consisting of a clinical examination, electrocardiogram (EKG) and echocardiogram.

The patient’s country of origin was recorded, and individuals were grouped as those from Central America (El Salvador, Guatemala, and Honduras) and those from Bolivia. We based our grouping on the general geographic distribution of *T. cruzi* discrete typing units[34, 35].

All patients had no symptoms of heart failure. Patients with no EKG or echocardiogram abnormalities and an ejection fraction (EF) greater than 50% were defined as non-cardiomyopathy patients. In Chagas disease, these patients are commonly referred to as being in the Indeterminate stage. While indeterminate-stage Chagas patients have a clear risk for the development of cardiomyopathy, the vast majority of these individuals will never develop any cardiac manifestations. The non-Chagas non-cardiomyopathy patients are likely to include individuals both with and without known risk factors for the development of cardiomyopathy, but a complete clinical workup was not performed. Patients with significant EKG abnormalities, many of which are known to be associated with Chagas disease, were defined as having asymptomatic early cardiomyopathy. These EKG changes were: RBBB, left bundle branch block (LBBB), LAFB, left posterior fascicular block (LPFB), any atrioventricular block, bradycardia of less than 50 beats per minute, non-specific intraventricular conduction delay (NIVCD), atrial fibrillation, atrial flutter, pathologic Q waves, PVCs, ventricular tachycardia, atrial pacing or ventricular pacing [12]. Electrocardiogram alterations per patient are described in **Supplementary Table 1**.

Patient serum was evaluated for anti-*T. cruzi* antibodies using the Hemagen Chagas kit (Hemagen Laboratories, Columbia, MD, USA), the Chagatest recombinant v.3.0 kit (Wiener Laboratories SAIC, Argentina), the Chagastest lysado (Wiener Laboratories SAIC, Argentina) and the IgG-TESA-blot. The IgG-TESA-blot was developed and performed using the trypomastigote excreted-secreted antigen from the *T. cruzi* Y strain [36, 37]. Individuals with positive results on two or more tests were considered seropositive for Chagas disease[32, 33]. 10 Chagas-positive patients were randomly selected, and 23 Chagas-negative patients were matched by age, sex, region of origin, and cardiac disease stage. The overall study design is summarized in Figure 1. The demographic and clinical features of the patients used for this study are summarized in Table 1.

**Table 1:** Study participant demographic data. P-value determined by Fisher’s exact test or student t-test

|  | Chagas-Positive |  |  | Chagas-Negative |  |  |
| --- | --- | --- | --- | --- | --- | --- |
|  | No evidence of cardiomyopathy (No CARD, Indeterminate) | Asymptomatic early cardiomyopathy (Early CCC) | p-value | No evidence of cardiomyopathy (No CARD) | Asymptomatic early cardiomyopathy (Early CARD) | p-value |
| Number | 4 | 6 |  | 13 | 10 |  |
| Mean Age (sd) | 51.85 (15.19) | 44.36 (4.59) | 0.279 | 47.22 (10.05) | 48.68 (12.25) | 0.756 |
| Female (%) | 3 ( 75.0) | 2 (33.3) | 0.524 | 9 (69.2) | 5 (50.0) | 0.417 |
| Bolivians (%) | 4 (100.0) | 4 (66.7) | 0.467 | 9 (69.2) | 6 (60.0) | 0.685 |
| Central Americans (%) | 2 (0) | 2(33.3) |  | 4(30.8) | 4(40.0) |  |

**Figure 1:**
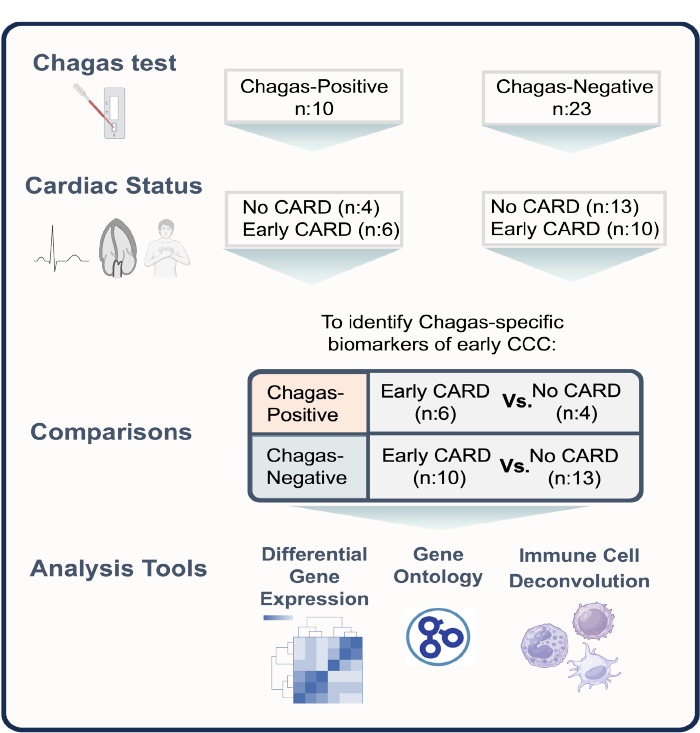
Study design schematic. 10 Chagas-positive patients were randomly selected, and 23 Chagas-negative patients were matched by age, sex, region of origin, and cardiac disease stage. These patients were stratified by Chagas status to compare cardiac patients with asymptomatic early cardiomyopathy (abnormal EKG, normal echocardiography, no symptoms of heart failure) to those with no signs or symptoms of cardiomyopathy using differential gene expression, gene ontology and immune cell deconvolution. Early CARD – asymptomatic early cardiomyopathy. Image created with BioRender.com

### Sample processing

Whole blood was obtained from patients and stored in DNA/RNA Shield (Zymo Research, Irvine, CA, USA). RNA was extracted using Quick-RNA Whole Blood Kit (Zymo Research, Irvine, CA, USA) and treated with TURBO DNAse (Thermo Fisher Scientific, Waltham, MA, USA) to remove remaining DNA contamination. Samples were then rRNA and globin depleted using the Ribo-Zero™ kit (Illumina, San Diego, CA, USA) and the GLOBINclear™ kit (Invitrogen, Waltham, MA, USA) respectively. Non-directional libraries were then constructed using Novogene’s proprietary kit and then loaded on the Illumina Novaseq 6000 S4 instrument per the manufacturer’s instructions. The samples were sequenced using a 2×150 Paired-End configuration. Paired reads were adapter- and quality-trimmed using Trimmomatic (v. 0.38), aligned to the human genome GRCh38 ensembl release 84 using HiSAT2 (v. 2.1.0), and raw read counts per gene were obtained using featureCounts (v 2.0.1). All other downstream analyses were conducted in R v.4.2.2.

## Differential expression analysis and quality checking

### Principal Component Analysis (PCA) and Differentially Expressed Gene (DEG) Identification

Raw counts were transformed using a variance stabilizing transformation (vst) to remove the dependence of the variance on the mean. PCA was performed on the transformed genes using base R stats prcomp and visualized using ggplot2 (v.3.4.0). Principal component analysis identified considerable variance introduced by batch effect and sex (Fig S1), so our analysis controlled for sequencing batch effects, and was stratified by sex.

Differential gene expression analysis was performed with DESeq2 [38], removing low-expressed genes which are defined as genes where less than 3 samples have at least 10 counts. P-values were determined using the Wald test and corrected for multiple testing using the Benjamini and Hochberg (BH) adjustment. Genes with a false discovery rate (FDR) of less than 0.1 and absolute fold change (|FC|) greater than 1.5 were considered significantly differentially expressed.

Heatmaps of the significantly (FDR < 0.1) differentially expressed genes were generated using the pheatmap package (v.1.0.12). Semi-supervised clustering of these genes (rows) was done using complete-linkage clustering of their vst transformed counts, and the distance between genes was determined by the Euclidian distance method.

### Gene Ontology Enrichment Analysis

Gene ontology enrichment analysis was performed using the enrichGO and gseGO functions of clusterProfiler (v. 4.4.4)[39, 40]. Both analyses were performed with a background gene set of all the genes submitted to DESeq2 analysis after filtering out low-expressed genes. enrichGO applies an over-representation analysis based on a one-sided Fisher’s exact test to the DEGs detected by DESeq2 analyses. gseGO applies a gene set enrichment analysis on the log2 fold change ranked list of background genes submitted to DESeq2 after filtering out low-expressed genes, based on the fgsea method[39, 41]. BH-adjusted p-values less than 0.05 are considered significantly enriched pathways. Gene set enrichment analyses were visualized using an enrichment plot. The enrichment plot was generated by using the pairwise_termsim function of clusterProfiler to calculate pairwise similarities of the enriched terms in a gene set using the Jaccard similarity index. This was then visualized with the emapplot function using a kk layout.

### Immune Cell Deconvolution

To obtain cell counts in transcripts per million (TPM), the previously quality trimmed fastq files were mapped and quantified with Salmon (v. 1.10.1) using a decoy-aware human transcriptome generated by concatenating the cDNA, ncRNA, and DNA primary assembly of the GRCh38 ensembl release 109. Quantification also corrected for fragment-level GC biases. Immune cell deconvolution of the TPM values was performed using ABSolute Immune Signal (ABIS) RNA-Seq deconvolution[42], CIBERSORTx with the LM22 signature-matrix and B mode batch correction[43], and xCell using the immunedeconv (v.2.1.0) package [44]. Groups were compared using the Wilcox-rank sum test with BH adjustment for multiple comparisons.

### Statistical analyses

Descriptive and inferential statistics were conducted using R v.4.2.2. Normality of data was assessed using visual examination and the Shapiro-Wilk test. Fisher’s exact test or student t-test were used where appropriate. Wilcox Rank sum test was used for other comparisons of continuous measures, with BH adjustment for multiple comparisons when more than two comparisons were evaluated.

## Results

### Transcriptional changes in peripheral blood are associated with early CCC progression

We sought to investigate whether gene expression changes in peripheral blood were associated with early heart disease and whether these signatures were different for the Chagas-positive and the Chagas-negative patients included in this study. We, therefore, compared individuals with no signs or symptoms of cardiac disease to those with asymptomatic cardiac abnormalities detectable by EKG in both Chagas-seropositive and seronegative individuals matched by age, sex, and region of origin (Table 1). Given the small sample size, we controlled for but did not stratify by sex.

In Chagas-positive patients, we identified 42 significant (FDR < 0.1, |FC| > 1.5) DEGs. Of these 21 were upregulated and 21 downregulated in early CCC compared to indeterminate stage individuals (Fig. 2a-b, Table S2). Notable downregulated genes reflect processes including: antigen presentation based on multiple major histocompatibility complex (MHC) classes (HLA-DRB1, HLA-DQB1, HLA-DOA), and CD1A[45]; T-cell activation (CD86) [46]; myeloid activation (CD300C)[47]; and activation of various immune cell types (nucleotide-binding oligomerization domain containing 2 (NOD2)) [48]. Upregulated immune genes reflect processes associated with both inhibitory and stimulatory signals in T cells and NK cells (killer cell lectin-like receptor B1 (KLRB1)) [49-51] and stimulatory roles in NK cells (killer cell lectin-like receptor C2 (KLRC2))[52].

**Figure 2.**
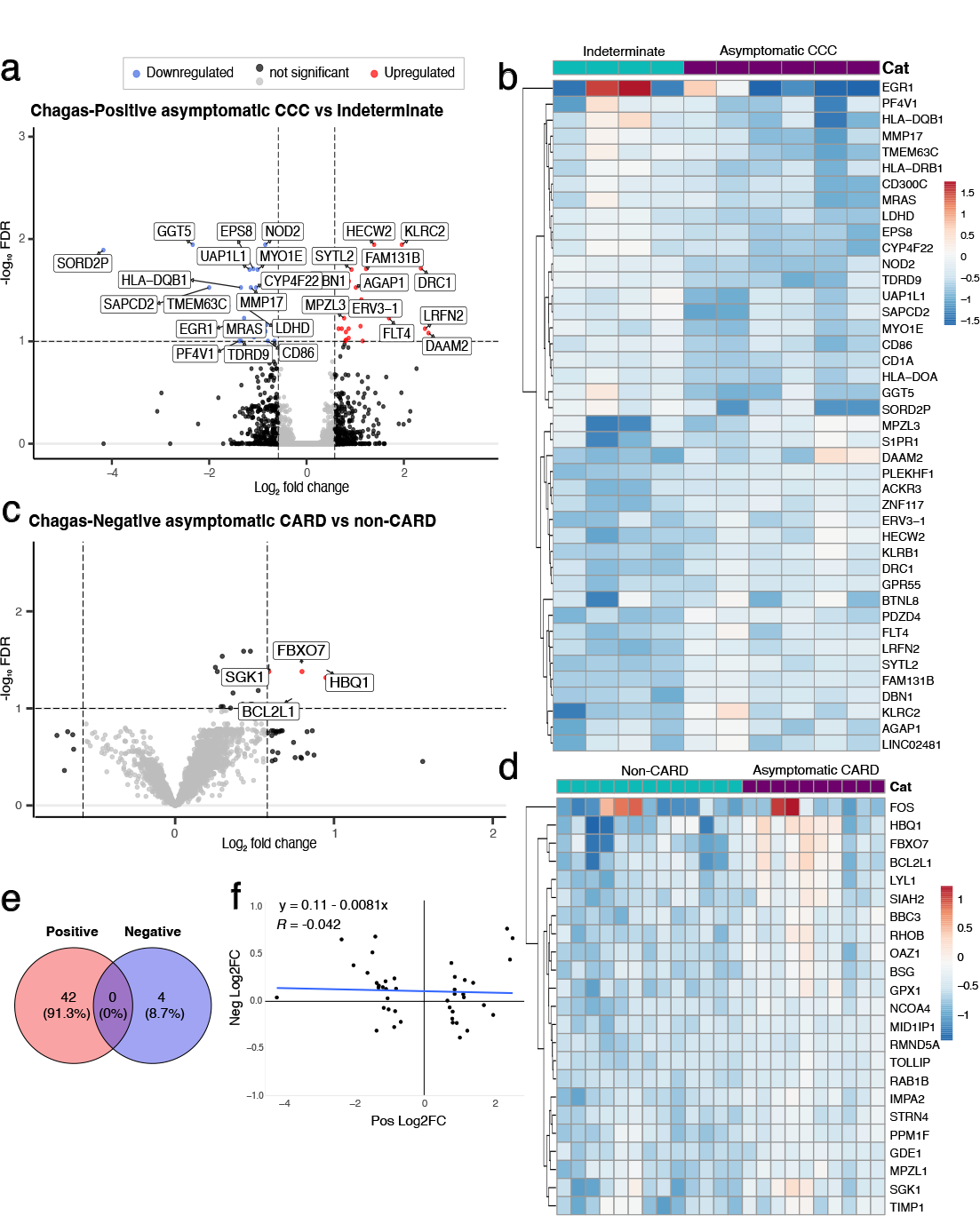
There are distinct gene expression changes that characterize early asymptomatic CCC. (a,c) Volcano plot of patients with asymptomatic cardiomyopathy vs those with no evidence of cardiomyopathy for a) chagas seropositive and c) seronegative individuals. Upregulated DEGs (FDR <0.1 and log2FC > 1.5) are shown in red, downregulated DEGs (FDR <0.1 and FC < - 1.5) are in blue, genes with FDR < 0.1 or |FC| > 1.5 but not both are in black, and non-significant DEGs are in grey. (b,d) Heatmap of relative vst-transformed values for the most significant DEGs with FDR < 0.05 in chagas b) seropositive and d) seronegative individuals. Color represents the sample’s difference from the mean for that gene. e) Venn diagram of significant DEGs of asymptomatic cardiomyopathy vs non-cardiomyopathy for chagas seropositive and seronegative individuals. f) scatter plot of log2FC of seropositive DEGs vs the log2FC of these same genes in seronegative individuals with Pearson correlation test. Trendline shown in blue. CARD- cardiomyopathy, CCC- chronic Chagas cardiomyopathy

Conversely, for seronegative individuals, we identified only 4 significant (FDR < 0.1, |FC| > 1.5) DEGs, all of which were upregulated in asymptomatic cardiomyopathy compared to patients with no evidence of cardiomyopathy (Fig 2c-d, Table S3). This smaller number of DEGs may be due to greater variability of etiologies and EKG alterations in seronegative individuals. Nevertheless, our data suggest that the underlying gene expression changes associated with early heart disease (asymptomatic cardiomyopathy vs. no evidence of cardiomyopathy) might be distinct in these Chagas-negative controls compared to Chagas-positive patients, as none of the significant DEGs intersect with those of Chagas-positive individuals (Fig. 2e). To ensure that this lack of intersection is not the result of small effect sizes in the non-Chagas individuals, we also performed a correlation analysis that showed no clear Association (R = -0.042, p = 0.79) between the effect sizes of the log_2_FC of the Chagas-positive DEGs and the log_2_FC of these same genes in Chagas-negative patients, which suggests that these DEGs are truly unique to early CCC pathogenesis (Fig 2f). Together these data indicate that dynamic changes in immune components accompany the development of early CCC.

### Early CCC progression is characterized by reduced antigen presentation and immune cell activation

Having demonstrated that there are early changes associated with CCC progression, we sought to explore the biologic processes underlying these changes. Using GO analysis, we found that in the Chagas group, no pathways were overrepresented in the upregulated genes, but 129 biologic processes were significantly over-represented (FDR <0.05) in the downregulated genes. These pathways were largely associated with antigen presentation and T cell function and were mainly driven by the DEGs: CD86, early growth response 1 (EGR1), HLA-DRB1, HLA-DQB1, HLA-DOA, NOD2, and CD1A (Fig 3a, Table S4). Immune cell deconvolution also demonstrates a trend towards a reduction in classical monocytes (antigen presenting cells) in Chagas early CCC compared to the indeterminate stage, though this finding did not reach statistical significance (Fig3b, S2). Conversely, T cell subsets, including mucosal-associated invariant T cells, CD naïve T cells, and CD8 memory T cells, as well as NK cells, trend towards an increase in Chagas stage B (Fig 3b, S2). This suggests that T cell activation is unlikely to be due to a reduction in T cell populations but may be due to fewer antigen presenting cells. These trends were not seen in Chagas-negative controls, suggesting that these immune cell alterations may be more indicative of CCC (Fig 3b,c). Furthermore, when analyzing non-Chagas disease patients with asymptomatic cardiomyopathy versus those with no evidence of cardiomyopathy, GO analysis showed no overlapping pathways with those of the Chagas disease group. These 24 biologic processes that were enriched in the non-Chagas disease asymptomatic cardiomyopathy group (compared to the non-cardiomyopathy group), were largely related to apoptotic mitochondrial processes and oxidative stress pathways (Fig4a, Table S4). These pathways have been strongly associated with ischemic, diabetic, and other causes of cardiomyopathy and heart failure[53]. This finding further highlights that CCC may have a distinct pathophysiology from at least some other forms of cardiomyopathy.

**Figure 3.**
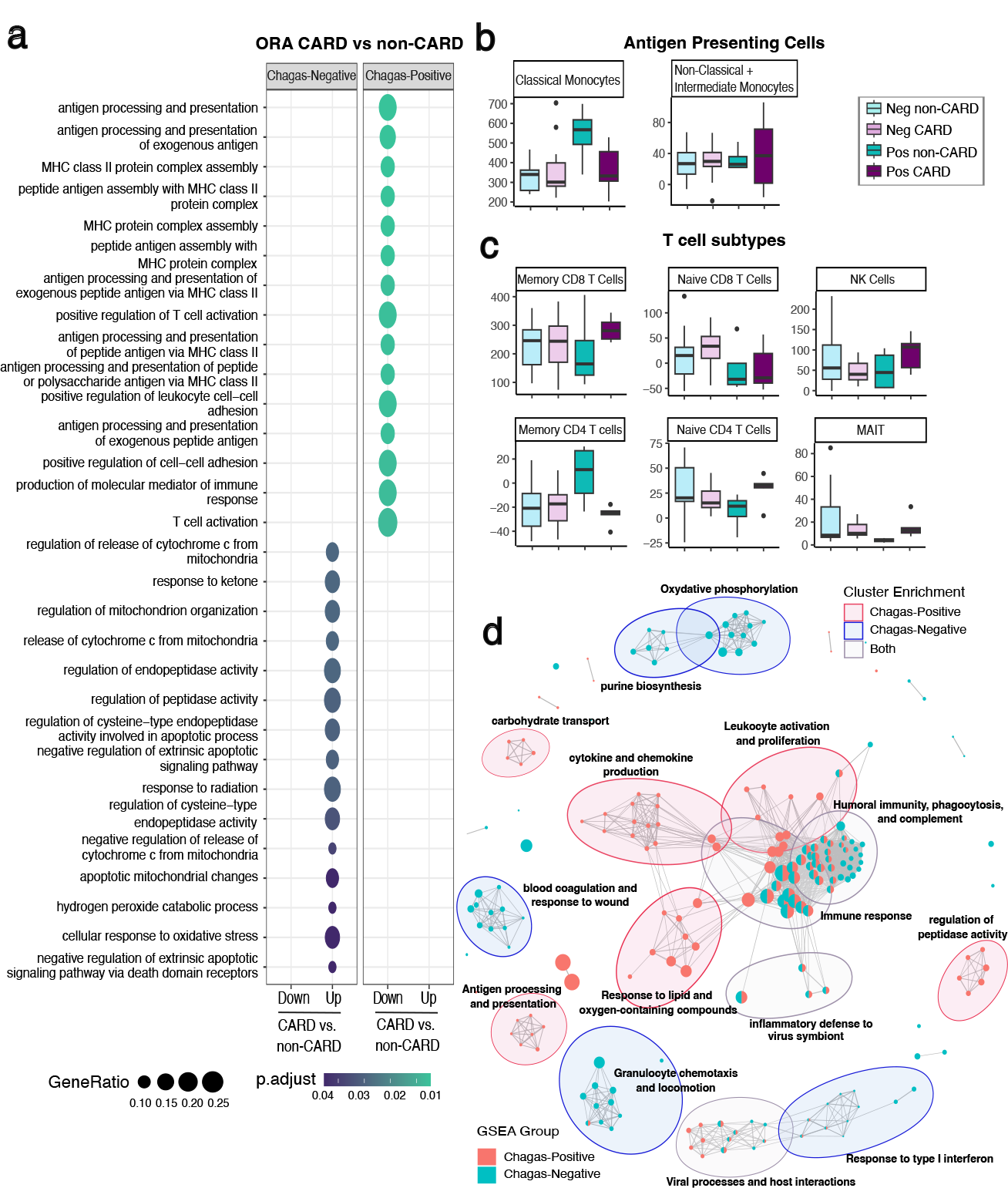
Asymptomatic stage B CCC demonstrates reduced antigen presentation and T cell activation. a) top 15 over-represented (ORA) GO biologic process (BP) terms (FDR <0.05) for downregulated (Down) and upregulated (Up) DEGs for asymptomatic cardiomyopathy vs non-cardiomyopathy patients that are Chagas seronegative (Neg) or seropositive (Pos). Circle size represents number of genes in the given pathway, color represents the FDR. b) box plot of antigen presenting cell deconvolution from bulk RNAseq data using ABsolute Immune Signal (ABIS) c) box plot of T cell deconvolution from bulk RNAseq data using ABIS d) enrichment map of gene set enriched analysis (GSEA) GO BP for seronegative and seropositive patients with lines connecting overlapping gene sets, and showing 15 clusters with > 2 biologic processes. Node size represents number of genes enriched in the pathway and line length represents pairwise similarity. Abbreviations: MAIT - Mucosal-associated invariant T cell, NK – Natural Killer, Pos – Chagas seropositive, Neg – Chagas seronegative, CARD-cardiomyopathy

To further verify the finding of CCC-specific immune alterations, we performed a gene set enrichment analysis of the complete fold change ranked gene list in order to detect gene changes in pathways that may be small and/or not statistically significant but occurring in the same coordinated direction. Here we identify 111 significantly (FDR <0.05) enriched biologic processes for the Chagas positive group, and 119 for the Chagas negative group (Fig 3d, Table S5), of which 46 are overlapping. While many of the overlapping processes are related to general aspects of immunity, there are notable immunological differences between the Chagas and non-Chagas groups. Chagas-positive individuals are enriched for pathways relating to cytokine and chemokine production, antigen presentation, leukocyte activation and proliferation, and response to oxygen and lipid-containing compounds, while Chagas-negative individuals are enriched for granulocyte chemotaxis, and response to type-I interferons. For non-immunologic processes, Chagas-positive individuals are enriched for pathways relating to the transport of carbohydrates, amino acids, and cholesterol, and regulation of peptidase activity. Conversely, Chagas-negative patients are enriched for coagulation, oxidative phosphorylation, and purine biosynthesis.

## Discussion

In this study, we have used peripheral blood samples from individuals with and without Chagas disease in different stages of heart failure to gain insight into the physiologic processes underlying progression to chronic Chagas cardiomyopathy. We demonstrate that, even though most chronic Chagas patients have ongoing chronic *T. cruzi* infections, patients developing early signs of CCC demonstrate immunologic alterations suggestive of a dysregulated immune response in the peripheral blood.

Notably, these changes are evident even before patients become clinically symptomatic and were not seen in patients with non-Chagas cardiomyopathies, suggesting that the mechanism by which CCC develops may be distinct from that of certain other non-Chagasic cardiomyopathies. These results highlight the important role that the immune system likely plays in CCC pathogenesis, while also demonstrating the feasibility of identifying patients in the earliest stage of CCC before they develop clinical symptoms.

Our results shed light on the immune mechanisms at play during the earliest stage of CCC. We found a notable downregulation of specific immune-associated genes in the peripheral blood of Chagas patients when comparing patients with asymptomatic electrocardiographic manifestations of Chagas disease to those in the indeterminate stage without any signs or symptoms of disease. In the early CCC patients, we observe a pattern of gene expression that strongly indicates a downregulation in antigen presentation and T cell activation in the peripheral blood. T cell activation is generally considered to depend on three signals 1) antigen presentation 2) co-stimulatory signaling and 3) cytokine signaling. In this study, we found a downregulation of genes corresponding to all three of these signals. First, various antigen-presenting genes were downregulated including various MHC molecules and CD1a which is capable of presenting non-peptide antigens to T cells [54]. For signal 2, the co-stimulatory protein CD86 was downregulated, and for signal 3, EGR1, an important transcription factor for various T cell stimulatory cytokines[55], was reduced. Consistent with our findings, Ferreira et al showed reduced HLA-DPB1 expression in moderate CCC compared to PCR-negative indeterminate stage patients[30] and *in vitro T. cruzi* infection has been shown to reduce expression of co-stimulatory molecules such as CD86[56] and antigen-presenting MHC proteins[57, 58]. In line with this, our immune cell deconvolution analysis suggests that although T cell numbers are not decreased, certain antigen presenting cells, which are required for T cell activation, may be decreased in the peripheral blood of early CCC patients. Together these data point toward a model in which T cell function is compromised in the peripheral blood of patients that develop early CCC, possibly due to a decrease in antigen presenting cells.

The mechanism by which compromised T cell activation may mediate cardiac damage in CCC has not been well explored. It has been shown that T cell terminal differentiation, exhaustion and depletion can result in cardiac inflammation and stress responses[59, 60]. It is also known that T cell responses are critical for controlling *T. cruzi* parasitemia based on knockout models of CD4, CD8, MHC-I, or MHC-II, which all show increased parasitemia and rapid host death upon *T. cruzi* infection [61, 62]. Thus, it is possible that an initial downregulation in anti-*T. cruzi* T cell responses may allow for greater parasite persistence and a low level of parasite-mediated cardiac damage, which after years of infection may ultimately lead to the electrocardiographic alterations seen in early CCC. It is then possible that once sufficient cardiac damage has occurred, immune responses may resurge due to the production of damage-associated molecular patterns and autoantigens[63]. This may then contribute to the largely pro-inflammatory findings that have traditionally been seen in studies of more advanced CCC[22, 26-29]. Most prior work has focused on severe CCC or has not distinguished between different stages of CCC for analysis. Such experimental designs, while certainly informative, are likely to mask transient changes specific to earlier stages of disease progression, and thus clouding our understanding of early CCC pathogenesis.

The current paradigm in the field is that CCC pathogenesis is largely a pro-inflammatory process [22, 26-29]. This logically stems from the infectious nature of Chagas disease, where one would expect a strong immune response against *T. cruzi* that could in turn cause tissue damage. While many studies have shown such pro-inflammatory patterns, there is a growing body of research on CCC showing changes similar to what we have observed in the current study, with various indicators of downregulated immune responses in CCC, such as downregulated innate immune markers, increased T cell exhaustion marker expression, and increased T regulatory cells[22, 64-67]. Indeed, a growing body of evidence in the field of cardiac immunology indicates that the immune system is intimately connected with cardiac function and dysfunction and that specific immune cell types can have disparate effects on the heart based on context[68]. Moreover, analyses of peripheral blood can reflect immune processes occurring locally in the cardiac tissue[69, 70], but this is not necessarily universally the case[71]. The immune downregulation we and others have observed, for instance, could reflect the loss of certain populations from circulation, as these immune cells home to other sites in response to infection and increasing cardiac damage. Thus, the complete picture of CCC pathogenesis is likely more nuanced, and potentially dynamic, than the current paradigm suggests. Our study is notable its focus on very early-stage CCC and highlights the value of stratifying patients by disease stage for comparison.

In addition, while there are limitations to the analysis of peripheral blood in CCC—most notably that it may not reflect the processes occurring at the site of cardiac damage—there is value in understanding the changes in this easily accessible tissue. We have shown here that we can find clear patterns of expression in the peripheral blood of asymptomatic early CCC patients. This suggests that it may be feasible in the future to identify blood biomarkers of disease progression even before the clinical EKG manifestations of disease. Longitudinal studies will be essential in identifying these markers of disease progression. For Chagas disease, it is currently impossible to predict or identify the 20-30% of patients who will progress to CCC. Anti-trypanosomal drugs are currently recommended for patients younger than 50 in the indeterminate stage of disease[72], but these drugs involve long treatment regimens and can be associated with notable side effects [8, 73]. As a result, it would be highly valuable to be able to risk-stratify the individuals that are most likely to develop CCC and that would most benefit from treatment. Peripheral blood biomarkers may thus become particularly important for informing treatment decisions in CCC and helping to identify those individuals in need of specialized treatment and closer follow-up. Indeed, it is possible that some of the differentially expressed genes identified in this study could be used in this way.

An additional important finding from this work is that the gene expression signature and immunologic processes associated with early CCC appear to be distinct in Chagas disease, as they were not seen in the other non-Chagas etiologies of early cardiomyopathy evaluated in this study. This comparison is limited by the small sample size of this study, which likely does not cover all the potential forms of non-Chagas cardiomyopathy, potentially masking meaningful differences that would be measurable with a more comprehensive control group. Nevertheless, while early CCC was associated with numerous downregulated immunologic processes—particularly antigen presentation and immune cell activation—non-Chagas early heart disease was more strongly associated with an upregulation in apoptotic mitochondrial processes and oxidative stress. These findings have important implications for the study and future management of CCC. If, as our results tentatively suggest, the underlying mechanisms mediating cardiac damage are distinct between Chagas and certain forms of non-Chagas heart disease, then the clinical management for these diseases should likely also be different. In particular, our findings suggest that immunomodulatory agents may prove useful for the management of early CCC. This may be particularly beneficial since anti-trypanosomal drugs are no longer effective at improving clinical outcomes once cardiac alterations have arisen[8].

Although this study provides important insight into the potential processes underlying early progression of CCC, it is subject to certain limitations. First, despite using early heart failure stages to understand early disease pathogenesis, this study is still limited by its cross-sectional design. Longitudinal analyses will be essential to truly understand the processes that precede disease progression and to understand immune dynamics throughout infection. In addition, our relatively small sample size, with large person-to-person variation, may have limited our ability to detect more subtle gene expression changes between groups. A larger sample size will also allow for a more thorough evaluation of sex, comorbidities, and *T. cruzi* strain differences in progression to early CCC, and will allow us to better compare Chagas progression to a wider variety of non-Chagas cardiomyopathies, with a more thorough evaluation of underlying risk factors. Furthermore, while differentially expressed genes are important in understanding the pathogenesis of CCC, we did not evaluate whether this reflects changes at the protein level; this will be necessary to fully understand the phenotypic changes in early CCC. Future studies will thus be needed to address these limitations.

Overall, we have identified key immunologic markers of *T. cruzi* infection and early chronic Chagas cardiomyopathy progression that help shed light on the pathogenesis of this disease. This work suggests that measurable changes in peripheral blood gene expression can be detected in CCC patients even before they become clinically symptomatic and that the mechanisms mediating progression to Chagas cardiomyopathy may be distinct from non-Chagas cardiomyopathies. These insights provide a path towards using peripheral biomarkers to identify patients most likely to progress to CCC, and also towards new immunologic treatment approaches for this early, and perhaps more treatable stage of disease.

## Acknowledgments

The authors thank the consulates of El Salvador, Mexico, Honduras, and Guatemala, and the community programs “Ventanilla de Salud,” Urban Strategies, and Family Networks in the Washington Metropolitan Area for facilitating access to Hispanic communities.

