## Supplemental Figure 1 and 2 for "Immunologic changes are detectable in the peripheral blood transcriptome of clinically asymptomatic Chagas cardiomyopathy patients"

### Supplemental Figures

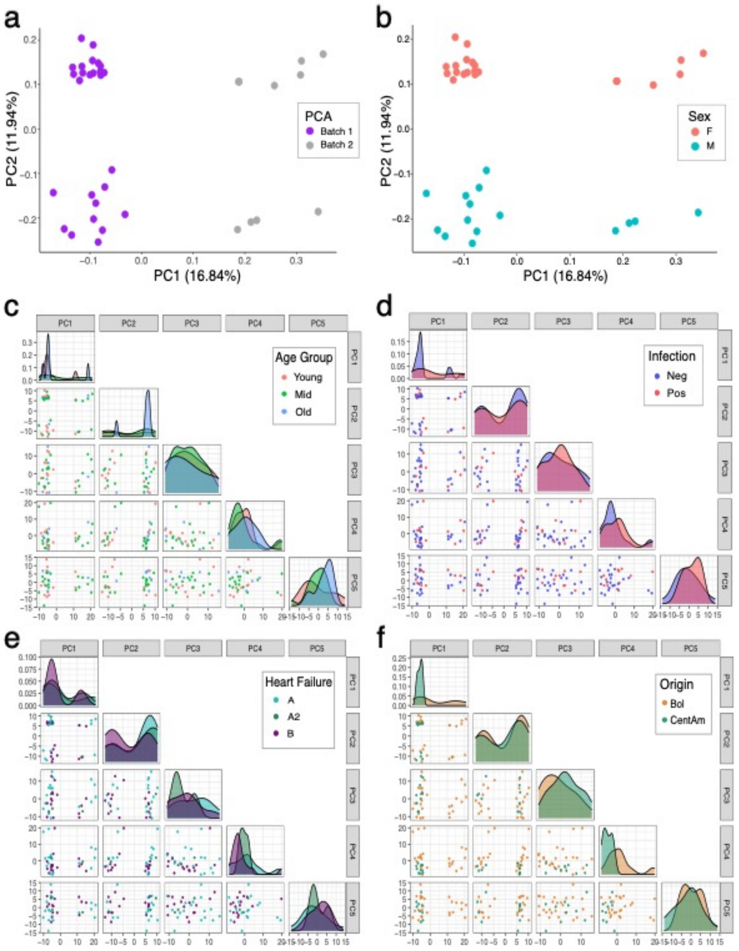


**Supplementary Figure 1. Sex and sequencing batch contribute to significant variance in the data.** Principal component analysis (PCA) plot of vst normalized gene counts. Principal component (PC) 1 vs PC2 colored by a) sequencing batch and b) sex. Pairwise plots of PC1 – PC5 with a density diagram along the diagonal and colored by c) age category, d) Chagas serostatus, e) heart failure stage, f) region of origin. F – female, M – Male, Neg – Chagas negative, Pos- Chagas Positive, Bol – Bolivian, CentAm – Central American.

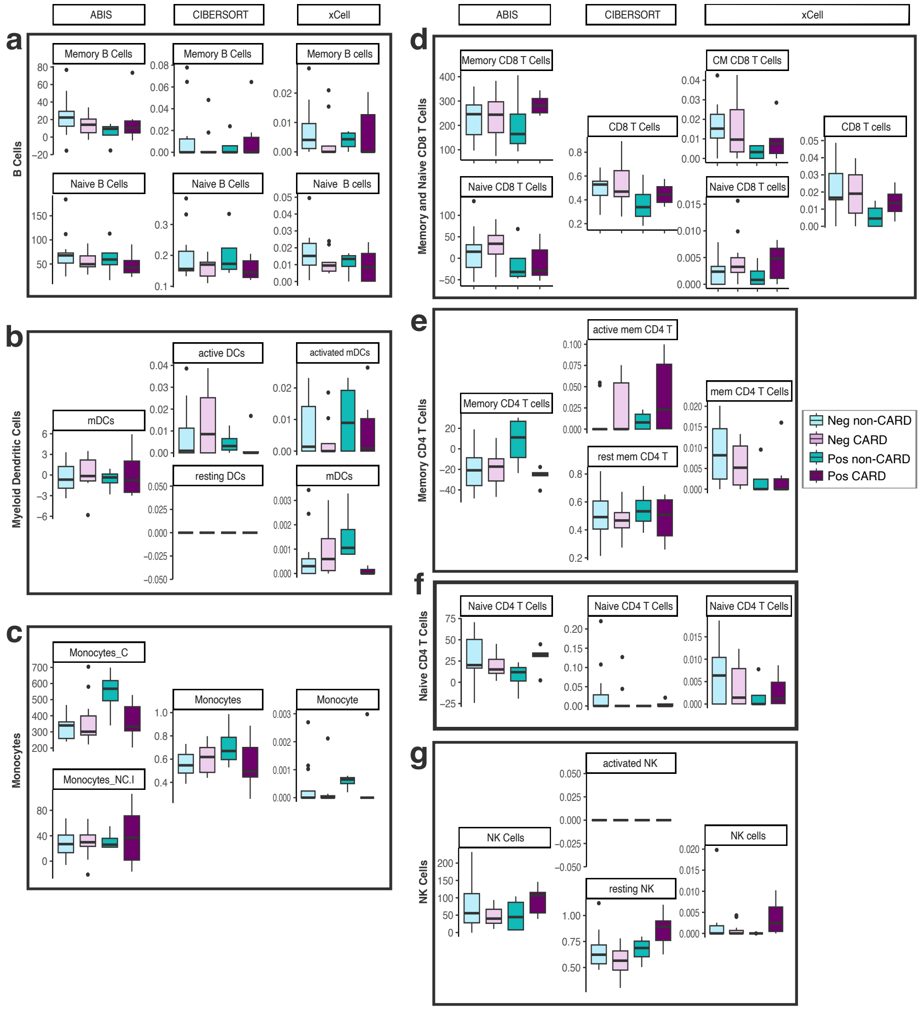


**Supplementary Figure 2: Antigen presenting cell and T cell subset changes are generally consistent across deconvolution methods.** box plot of B cell deconvolution of bulk RNAseq data for Chagas-negative stage A, Chagas-negative stage B, Chagas-positive stage A and Chagas-positive stage B patients. Deconvolution was performed using ABsolute Immune Signal (ABIS), CIBSERSORTx and xCell for a) B cells, b) myeloid dendtritic cells, c) monocytes d) CD8 T cells, e) Memory CD4 T cells f) Naïve CD4 T cells, and g) Natural Killer cells. Abbreviations: C – Classical, NC + I – Non-Classical and Intermediate, CM – Central memory, Mem – memory, NK – Natural Killer, Pos – Chagas seropositive, Neg – Chagas seronegative
